# The mutational landscape of single neurons and oligodendrocytes reveals evidence of inflammation-associated DNA damage in multiple sclerosis

**DOI:** 10.1101/2022.04.30.490132

**Authors:** Allan Motyer, Stacey Jackson, Bicheng Yang, Ivon Harliwong, Wei Tian, Wingin Shiu, Yunchang Shao, Bo Wang, Catriona McLean, Michael Barnett, Trevor J. Kilpatrick, Stephen Leslie, Justin P. Rubio

**Affiliations:** Melbourne Integrative Genomics, The University of Melbourne, Victoria 3010, Australia; School of Mathematics and Statistics, The University of Melbourne, Victoria 3010, Australia; School of BioSciences, The University of Melbourne, Victoria 3010, Australia; The Florey Institute of Neuroscience and Mental Health, Victoria 3052, Australia; MGI-Australia, 300 Herston Rd, Herston, Queensland 4006, Australia; BGI-Australia, 300 Herston Rd, Herston, Queensland 4006, Australia; China National GeneBank, Jinsha Road, Shenzhen 518120, China; State Key Laboratory of Quality Research in Chinese Medicine, Institute of Chinese Medical Sciences, University of Macau, Macao 999078, China; Victorian Melanoma Service, Alfred Health, Melbourne, Victoria 3004, Australia; Brain and Mind Centre, The University of Sydney, NSW 2006, Australia; Sydney Neuroimaging Analysis Centre, Camperdown, NSW 2050, Australia; Florey Department of Neuroscience and Mental Health, The University of Melbourne, Victoria 3010, Australia; Department of Neurology, Royal Melbourne Hospital, Victoria 3050, Australia

## Abstract

Neuroinflammation has been linked to DNA damage in multiple sclerosis (MS), but its impact on neural cell genomes at nucleotide resolution is unknown. To address this question, we performed single nucleus whole genome sequencing to determine the landscape of somatic mutation in 172 neurons and oligodendrocytes (OLs) extracted from post-mortem brain tissue from 5 MS cases and three controls. We identified two cases with a significant excess of somatic single nucleotide variants (sSNV) in neurons and OLs from MS inflammatory demyelinated lesions. For a case with primary progressive MS, this translated to a 68% increase in sSNV frequency and 32-year equivalent increase in biological age of lesion-resident cells. Mutational signature analysis conducted on all cells revealed that defective DNA repair and transcription-associated DNA damage are important mutagenic mechanism in both neurons and OLs in MS. Our findings provide the first evidence that inflammation in the brains of people with MS is associated with DNA damage, which may have implications for other neurodegenerative diseases and future drug development.

## Introduction

Multiple sclerosis (MS) is the most common inflammatory disease of the central nervous system, affecting 2-3 million people globally (*1, 2*). MS generally starts early in adult life, and often leads to severe disability until death, most often in the sixth to seventh decade (*1*).

The clinical symptoms of MS arise when autoreactive immune cells infiltrate the brain and cause injury to resident cells. Oligodendrocytes (OLs), a type of glial cell in the CNS that myelinates axons and facilitates the transmission of action potentials, are the target of immune insult in MS, with ensuing damage setting in motion a cascade of events that results in axonal injury and neurodegeneration (*3, 4*). Thus, neuroinflammation and neurodegeneration are causally linked in MS, but our understanding of the intervening pathophysiological mechanisms is incomplete (*5, 6*). In MS, the natural process of myelin repair becomes inadequate over time, hence regenerative therapies that may be used in combination with existing disease modifying immunotherapies remain an attractive prospect (*7*). Consequently, it follows that understanding processes linking neuroinflammation and neurodegeneration in MS is crucial to the overall goals of protection against damage and also regeneration.

Damage to the nuclear and mitochondrial DNA of neurons has been speculated to drive natural aging as well as neurodegenerative disease (*8, 9*). For neurodegenerative diseases such as motor neurone disease (MND), the argument for a pivotal role is strong, given the involvement of MND genes in DNA damage and its repair (*10-13*). For MS, our knowledge of the potential for DNA damage being a factor in neurodegeneration is limited to observations of increased amounts of oxidized guanine (8-oxoG) in both neurons and OLs in inflammatory demyelinated brain lesions post-mortem (*14-16*). However, while suggestive, it is unclear whether these findings are reflective of a chronic process, such as neuroinflammation extending for years or decades, and if the changes observed would ultimately translate to a permanent impact on genome integrity.

Single nucleus whole genome sequencing (snWGS) studies of neurons conducted to date have shown that somatic single nucleotide variants (sSNVs), which equate with DNA damage, accumulate at a rate of 25 sSNVs per annum in healthy individuals (*17-19*), reaching on average 2000-3000 sSNVs in individuals aged ∼80 years and being more abundant in individuals with the rare progeroid diseases, Cockayne syndrome and xeroderma pigmentosa (*19*).

To determine whether neuroinflammation could be driving somatic mutagenesis in MS, we have conducted the largest snWGS study for a common neurological disease to date, the first such study for MS, and the first investigation of the mutational landscape of OLs.

## Methods summary

We have developed a novel method for isolating single OL nuclei from post-mortem MS and control brain tissue, whilst also isolating neuronal nuclei from the same sample (**Fig. 1**). A key aspect of our study design is that for each MS case we investigated single OLs and neurons from both pathology-affected (regenerated and/or chronic lesion) and pathology-free, normal appearing tissue samples, while also doing the same for age-matched controls. We have then conducted snWGS to identify sSNVs, followed by statistical and computational analysis to derive mutational signatures and determine potential differences between cell types from pathology-affected and pathology-free tissue.

**Fig. 1.**
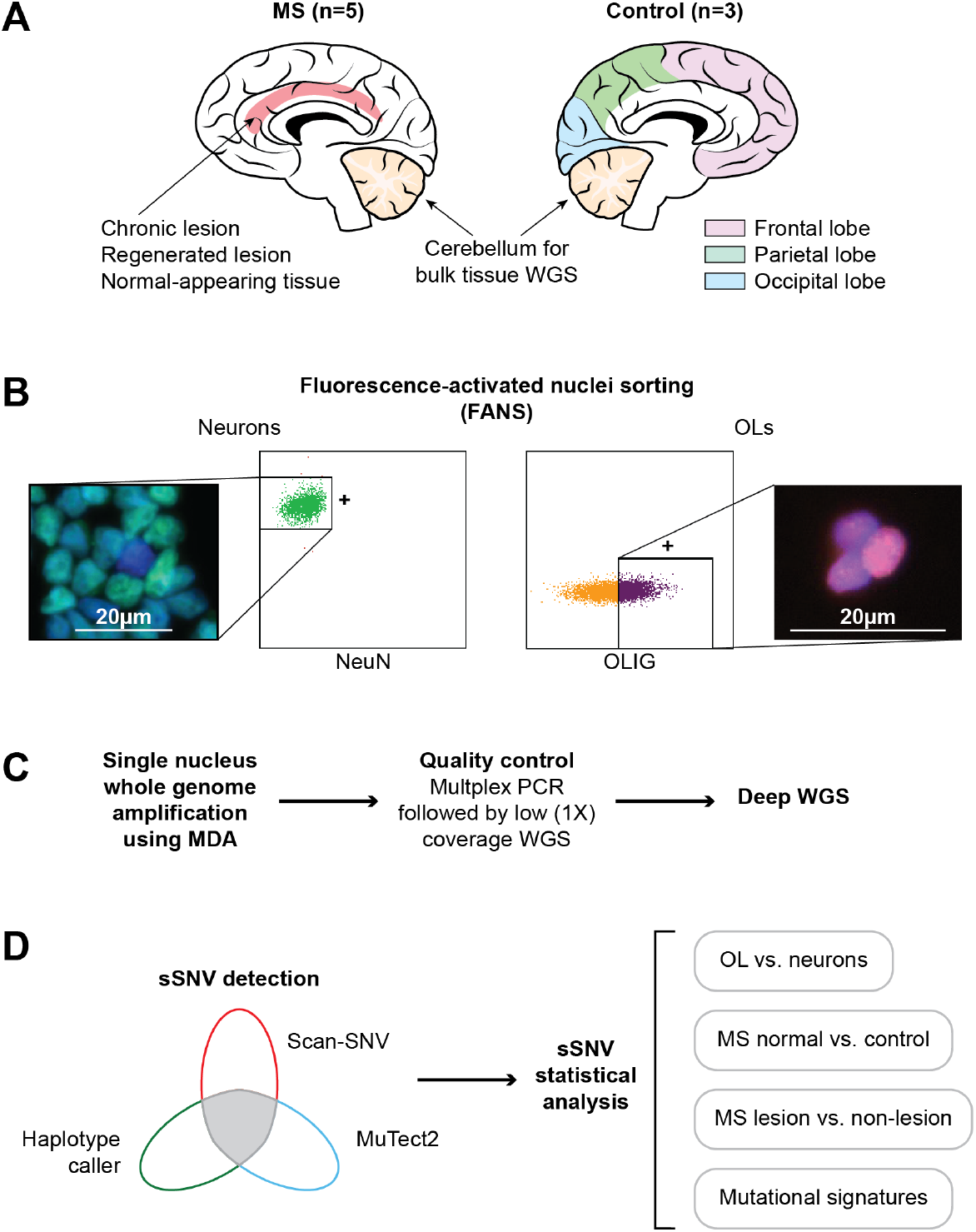
Experimental approach to determine whether somatic mutations are associated with MS pathology. **(A)** Frozen neuropathological tissue was obtained from five MS cases – pathology-affected (chronic and regenerated lesions) and pathology-free (normal-appearing) –and three controls – frontal lobe (Fr lobe), parietal lobe (Par lobe) and occipital (Occ lobe). Genomic (g)DNA from cerebellum (bulk) tissue was subjected to deep WGS and used to detect germline SNVs. **(B)** Other MS and control tissues were processed (see methods) and subjected to fluorescence-activated nuclei sorting (FANS), to isolate neuronal and OL nuclei using NeuN and OLIG transcription factors markers. Fluorescent microscopy images show NeuN+ and OL+ sorted nuclei co-stained with DAPI. **(C)** Cold alkaline lysis (*21*) of single nuclei was performed prior to whole genome amplification using multiple displacement amplification (MDA) (*22*). The first quality control (QC) step consisted of multiplex PCR (*18*) and, for the subset of snWGA reactions passing this step, low coverage (∼1X) WGS was performed to identify the highest quality samples for deep WGS. **(D)** A triple-called approach, using SCAN-SNV (*23*), Mutect2 (*24*) and HaplotypeCaller (*25*), was used to identify sSNVs for statistical analyses based on cell type, tissue type and disease status.

### Post-mortem brain tissue

MS neuropathological tissue for five MS cases was obtained from the MS Research Australia Brain Bank (**Table 1**). Lesional tissue was characterized microscopically as showing evidence of immune-mediated damage (neuroinflammation) and for molecular evidence of remyelination, with regenerated lesions showing evidence of myelin repair and chronic lesions showing a lack of myelin repair (refer to Supp Methods). OLs and neurons extracted from these lesional tissue samples were selected as those likely to have been exposed to immune insult above and beyond those from pathology-free, MS normal tissue. Independent control tissue was obtained post-mortem from the Victorian Brain Bank for three individuals who died from non-neurological disease causes. Given previously observed age-related and regional differences in sSNV abundance in neurons (*19*), controls were selected spanning the age range of MS cases and tissue samples (frontal, parietal and occipital lobes) approximately matched to regions of the brain from where lesional and MS normal tissue samples were taken (**Table 1**).

**Table 1.** MS case and control information. **A**. Information for individual MS cases and controls from which neuropathological tissue samples were obtained. **B**. Number of single cell samples analyzed, by individual, cell type (neuron or OL), tissue pathology (for MS cases), and brain region (for controls).

| Case ID | Age-at-death | Sex | MS subtype | PMI (hrs) | Disease duration (yrs) | Received MS DMI | Cause of death |
| --- | --- | --- | --- | --- | --- | --- | --- |
| MS1 | 68 | F | SPMS | 15 | 33.5 | Yes | HBI, UAO, MS |
| MS2 | 69 | F | SPMS | 8.5 | 38 | N/A | APa, MS |
| MS3 | 72 | F | SPMS | 4 | 38.1 | No | Pa, MS |
| MS4 | 73 | M | PPMS | 25.5 | 15.4 | No | Ge, PVD, IHD, MS |
| MS5 | 68 | F | SPMS | 7 | 26.7 | Yes | Rs, Hs, RF, AMI, MS |
| Control ID |  |  |  |  |  |  |  |
| CT1 | 52.1 | M | - | 33 | - |  | HS |
| CT2 | 75.9 | M | - | 50 | - |  | AMI |
| CT3 | 63.9 | M | - | 68 | - |  | IHD |

**B**
| Case ID | Normal (Neu/OL) | Regenerated Lesion (Neu/OL) | Chronic Lesion (Neu/OL) | Total All Tissues (Neu/OL) |
| --- | --- | --- | --- | --- |
| MS1 | 4 / 3 | 0 / 0 | 4 / 3 | 8 / 6 |
| MS2 | 5 / 6 | 0 / 6 | 6 / 6 | 11 / 18 |
| MS3 | 6 / 6 | 0 / 0 | 6 / 3 | 12 / 9 |
| MS4 | 6 / 6 | 6 / 6 | 4 / 6 | 16 / 18 |
| MS5 | 5 / 6 | 6 / 6 | 3 / 6 | 14 / 18 |
| Total MS | 26 / 27 | 12 / 18 | 23 / 24 | 61 / 69 |
| Control ID | Frontal cortex (Neu/OL) | Occipital cortex (Neu/OL) | Parietal cortex (Neu/OL) | Total All Tissues (Neu/OL) |
| CT1 | 1 / 4 | 3 / 2 | 3 / 2 | 7 / 8 |
| CT2 | 2 / 2 | 0 / 3 | 3 / 3 | 5 / 8 |
| CT3 | 3 / 4 | 2 / 2 | 1 / 2 | 6 / 8 |
| Total Controls | 6 / 10 | 5 / 7 | 7 / 7 | 18 / 24 |

**Abbreviations:** Secondary progressive (SPMS), primary progressive MS (PPMS). Hypoxic brain injury (HBI), upper airway obstruction (UAO), aspiration pneumonia (APa), pneumonia (Pa), gangrene (Ge), peripheral vascular disease (PVD), ischemic heart disease (IHD), rhabdomyolysis (Rs), heatstroke (Hs), renal failure (RF), acute myocardial infarction (AMI), hemorrhagic shock (HS).
**Clinical note (Table 1):** In about half of people with the relapsing remitting (RR) form of MS, sporadic bouts of immunological activation (NI) lead to a neurodegenerative, secondary progressive (SP) disease phenotype 10-20 years after onset. In 5-10% of cases, disease starts with a primary progressive (PP) phenotype that displays no overt clinical signs of immunological activation, but the hallmarks of NI can still be observed in post-mortem brain tissue as demyelinated lesions by a neuropathologist and via magnetic resonance imaging.

### Fluorescence-activated nuclei sorting (FANS)

This technique was used to isolate OL and neuronal nuclei from frozen post-mortem brain tissue, using a published method for neurons (*20*), with some modifications, and a new method we developed for OLs using antibodies against OLIG transcription factors (Supp Methods, **Fig. S1** and **Fig. S2**). Care was taken to avoid systematic errors in sample processing by ensuring that flow cytometry was standardized across samples, with tissue samples for every case or control processed and run on the same day on the same machine by the same operator with randomized running order of lesional and MS normal tissue samples. Further, flow sorting of OL and neuronal nuclei, processed from the same tissue preparation split 50:50, occurred on consecutive flow sorts, with the order switched for each tissue sample.

### Single nucleus whole genome sequencing (snWGS) and somatic mutation detection

Cold alkaline lysis of flow sorted nuclei (*21*) was performed prior to whole genome amplification (WGA) using multiple displacement amplification (MDA) (*22*). To identify the highest quality samples, we initially performed multiplex PCR on all single nuclei WGA reactions (*18*), followed by low coverage (∼1X) snWGS on the subset that passed PCR quality control (**Fig. 1** and Supp Methods). Samples passing both quality controls steps were selected for deeper coverage snWGS.

Overall, we conducted snWGS at 66X mean coverage on 172 neuronal and OL nuclei from MS cases (n=130) and controls (n=42) (**Table 1B**; **Table S1**). Bulk tissue genomic DNA samples were extracted from cerebellar tissue of cases and controls and were sequenced to 76.5X mean coverage. sSNVs in single cells were called from the consensus output of three variant calling programs: SCAN-SNV (*23*), GATK Mutect2 (*24*) and GATK HaplotypeCaller (*25*) and leveraging bulk tissue WGS data for calling germline SNVs. Across all 172 single cells, 155,021 triple-called sSNVs were detected (**Table S2**).

## Results

### Somatic mutation burden

For both neurons and OLs in MS normal and control tissue, we observed on average 1831 sSNVs per cell (n=95), with a large range of sSNV counts, 860-4112, for individual cells. We observed trends of excess sSNVs in neurons compared to OLs in both control (11.2%, 95% CI (−2.9%, 27.3%), p=0.13, n=42) and MS normal (10.3%, 95% CI (−3.4%, 25.4%), p=0.15, n=53) tissue, which was statistically significant when cells were combined across tissues (10.9%, 95% CI (1.0%, 21.7%), p=0.03, n=95) (**Fig. 2A**). This difference equated to a mean excess of 184 sSNVs in neurons compared to OLs. A non-statistically significant difference between sSNVs per cell in MS normal versus control tissue for both cell types was also observed (5.5% excess in MS normal tissue, 95% CI (−16.1%, 32.8%), p=0.70, n=95) (**Fig. 2B**). A statistically significant association between sSNV count and age was observed (1.66% increase per year, 95% CI (0.78%, 2.51%), p=0.008, n=172), which translated to an increase of ∼30 sSNVs per year (**Fig. 2C**). This rate of sSNV increase per annum is similar to that observed previously in single neurons (*19*), giving us confidence in our protocols and analysis.

**Fig. 2.**
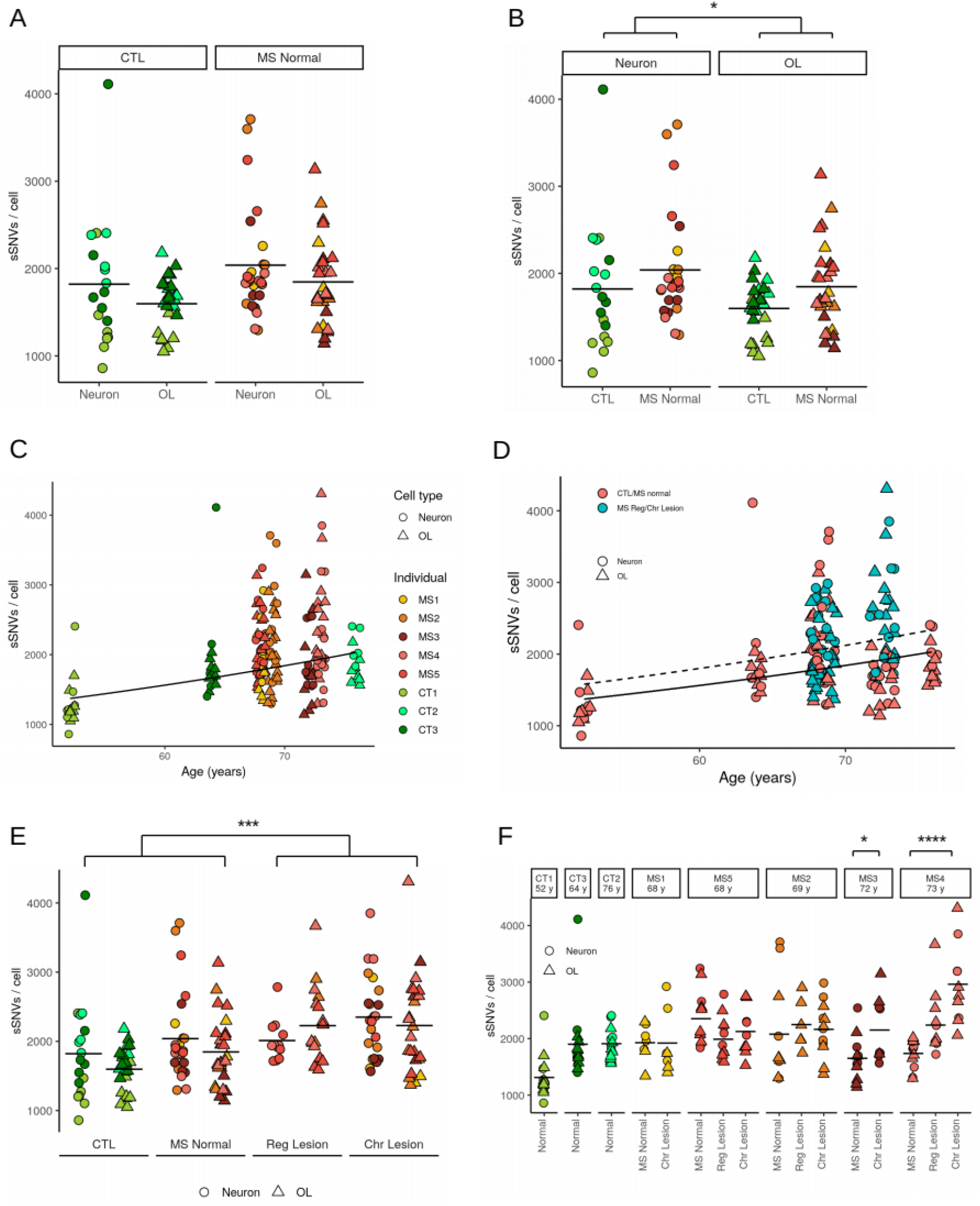
sSNV frequencies differ by cell type, age and tissue pathology. **(A & B)** To visualize sSNV counts in neurons and OLs from control (CTL) tissue and normal-appearing MS tissue (MS normal), respectively, two plots illustrate the mean number of counts for each comparison denoted by a horizontal line. Statistical analysis revealed sSNVs are increased modestly in neurons vs. OLs when counts from CTL and MS normal were combined. A non-significant trend of increased sSNVs is observed for both cell types in MS normal vs. CTL. **(C)** Linear regression of age vs. sSNV counts for MS cases and controls revealed an age-associated increase, which is similar to that previously described for neurons alone (*19*). **(D and E)** sSNV counts are significantly increased in neurons and OLs from MS lesion (dashed linear regression line) vs. those from non-lesion (solid regression line) tissue. **(F)** sSNV counts viewed in each MS case shows that the difference between MS lesion and non-lesional tissue is driven by two MS cases (MS3 and MS4). Significance levels indicated by asterisks: * p < 0.05, *** p < 0.001, **** p < 10^−5^.

Separation of neuronal and OL samples based on tissue of origin revealed a statistically significant sSNV excess in cells from MS lesion compared to MS normal and control tissue (mean 15.2% increase (95% CI 6.6%, 25.7%), p=0.0008, n=172) (**Fig. 2 D, E**). Analyses accounted for non-independence of cells from the same individual and age-associations (see Supp Methods).

To determine whether the sSNV excess observed was consistent across MS cases, we considered the data for individuals separately, and observed that two of the five MS cases (MS3 and MS4) were driving the association between sSNV count in lesion vs. MS normal tissue (**Fig. 2F**). The strongest association was observed in MS4 (27.4% excess in regenerated lesion vs MS normal, 95% CI (8.0%, 50.3%); 68.2% excess in chronic lesion vs MS normal, 95% CI (41.4%, 100.0%); p=5×10^−6^, n=34), a 73-year-old male with primary progressive (PP) MS who had the shortest disease duration (15.4 years) and who was negative for HLA-DRB1*15:01, the strongest genetic risk factor for MS (*26*) (**Table 1**). MS3, an HLA-DRB1*15:01 heterozygous 72-year old female with secondary progressive (SP) MS, had a longer disease duration (38.1 years) (29.6% excess in chronic lesion vs non-lesion, 95% CI (3.7%, 61.8%), p=0.02, n=21). No statistically significant relationship was found between sSNV count and tissue type in MS1 (p=0.85, n=14), MS2 (p=0.63, n=29) or MS5 (p=0.15, n=32).

Putting these results into context, the increase in OL and neuronal sSNVs from chronic lesion vs MS normal tissue is equivalent to 31.6 years of cellular aging for MS4 and 15.7 years for MS3, respectively.

### Mutational signatures

Pioneering studies conducted in the cancer field have demonstrated that somatic mutagenesis can be influenced by both exogeneous (e.g. tobacco) and endogeneous (e.g. transcription) factors, and that they can leave a detectable imprint or signature in the genomic sequence (*27, 28*). Therefore, to determine the mutational processes impacting OLs and neurons in the MS brain, we conducted mutational signature analysis (*27, 29*).

Across all 172 MS and control single OL and neuron samples, four statistically robust mutational signatures (MS_Sig1-Sig4) – each signature consisting of different relative contributions of six possible mutation types within their 16 trinucleotide contexts – and their contributions to the sSNVs of each sample were inferred (**Fig. 3A, Fig S3**). MS_Sig1-Sig4 have varying degrees of similarity to previously observed signatures in the Catalogue Of Somatic Mutations In Cancer (COSMIC) database (*30*) (**Fig. 3B**). From an aetiological perspective, it is noteworthy that MS_Sig1 is most similar to SBS44, a signature linked to defective DNA mismatch repair (*31*), while MS_Sig2 is most similar to SBS30, a signature linked to deficiency in base excision repair (*32*). MS_Sig3 is most similar to SBS19, a signature of unknown aetiology in cancer and MS_Sig4 is most similar to SBS5, an age-associated signature observed across both health and disease in humans and other mammalian species (*32, 33*).

**Fig. 3.**
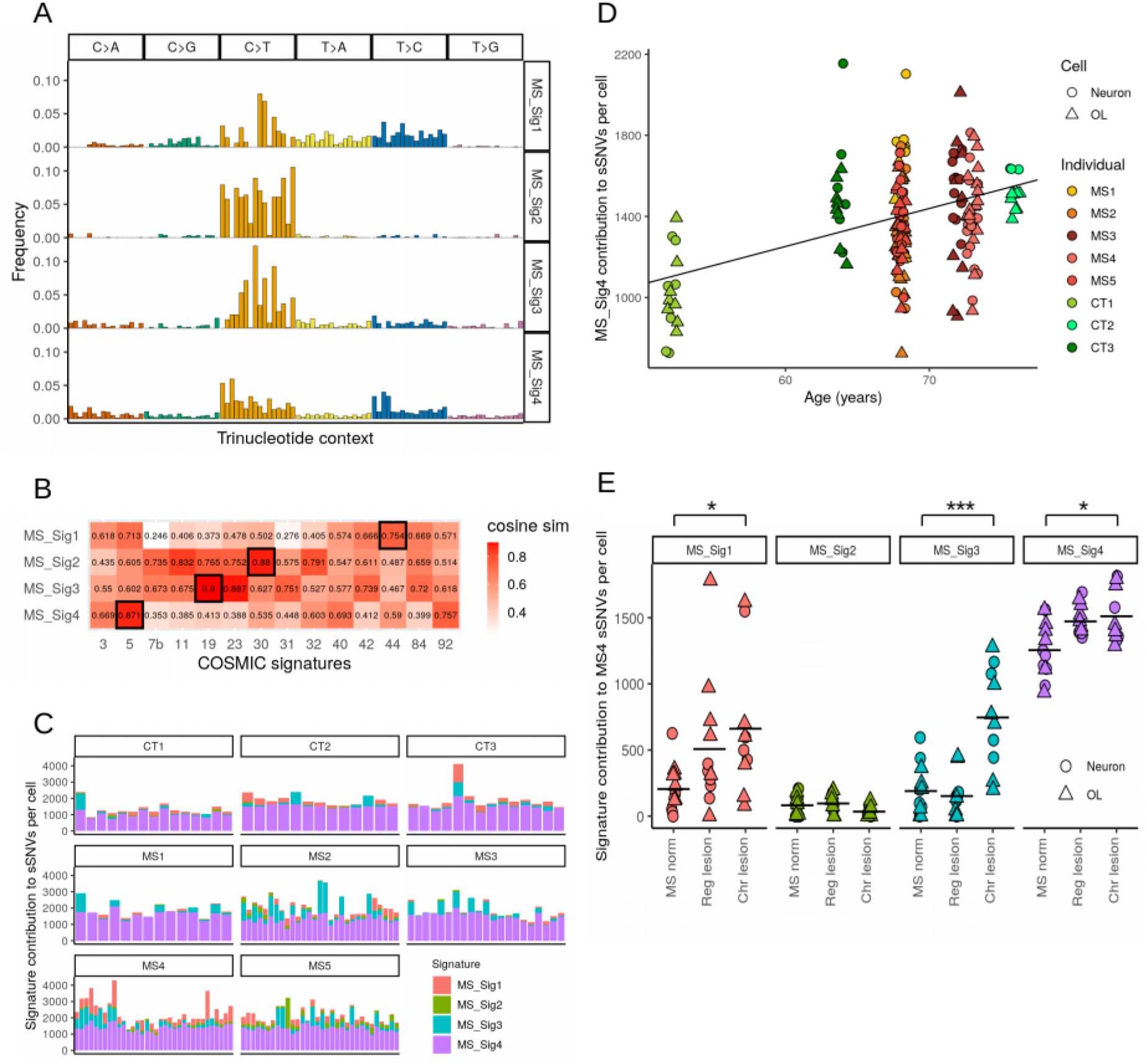
Mutational signatures. **(A)** Four mutational signatures were identified from sSNVs in 172 neurons and OLs using SignatureAnalyzer (*29*). Each signature specifies a probability distribution over six possible mutation types and their 16 possible trinucleotide contexts. **(B)** Cosine similarities between derived mutational signatures and the most similar signatures in the COSMIC database. **(C)** SignatureAnalyzer was used to infer the contribution of each of the four mutational signatures to the total sSNV count of single OLs and neurons from cases and controls, each represented by a bar on the x-axis **(D)** Plot showing contribution of MS_Sig4 to sSNV count is correlated with age for MS cases and controls. **(E)** Contribution of mutational signatures to sSNV counts in different tissues from MS4, with statistically significant increases in signature contributions for chronic lesion tissue vs. normal appearing tissue for MS_Sig1, MS_Sig3 and MS_Sig4. Significance levels indicated by asterisks: * p < 0.05, *** p < 10^−4^.

Next, we determined the contribution of the four mutational signatures to sSNV counts in all 172 cells. We observed that MS_Sig4 dominated sSNV counts in all single OLs and neurons (**Fig 3C, Fig S4**) and found it to be positively associated with age (18.7 sSNVs per year, 95% CI (5.8, 31.7), p=0.032, n=172) (**Fig 3D**). Given its age association, and the fact that none of the other three signatures were associated with age in either cell type from pathology-free or MS lesional tissue (**Fig S5**), MS_Sig4 is likely equivalent to signature A, as identified recently in single neurons (*19*). Finally, MS_Sig4 was also the only signature that contributed, at a level of statistical significance, to the difference in sSNV count between neurons and OLs in pathology-free tissue (mean 90.4 increase in neurons, 95% CI (3.4, 177.6), p=0.044, n=95, age included as covariate) (**Fig S6**).

Given the association between sSNV counts and MS tissue type was heterogeneous across MS cases, we investigated mutational signature contributions to sSNV counts at the individual level (**Fig 3E, Fig S7**). For MS4, the case with the strongest association between sSNV count and tissue type, we observed increased signature contributions for chronic lesion tissue vs. normal appearing tissue for MS_Sig1 (mean increase 456.0 sSNVs, 95% CI (93.4, 818.6), p=0.015), MS_Sig3 (mean increase 555.9 sSNVs, 95% CI (336.9, 774.9), p=1.5×10^−5^) and MS_Sig4 (mean increase 255.8 sSNVs, 95% CI (98.8, 412.9), p=0.0023). In MS3, the other case with excess sSNVs in lesional OLs and neurons, only MS_Sig4 had a statistically significant increase in its contribution to sSNV counts in lesion tissue (mean increase 330.5 sSNVs, 95% CI (121.0, 539.9), p=0.0037). Conversely, for MS3 there was a statistically significant decrease in the contribution for MS_Sig1 in lesion tissue (mean decrease 85.6 sSNVs, 95% CI (31.0, 140.1), p=0.0039), counter to the overall increase in lesion tissue, highlighting the heterogeneity of cases. Statistically significant differences in signature contributions were not observed for other signatures and MS cases.

Taken together, these results highlight the heterogeneous mutational processes underlying DNA damage and its repair in the MS brain, which could be underpinned by both variability in the type and quantum of mutagenic factors generated during inflammatory demyelination (e.g. reactive oxygen species), combined with deficiency in endogenous pathways of DNA repair at the individual level.

### Transcriptional strand asymmetry

Differential efficiency of DNA repair mechanisms operating on the transcribed strand (TS) and non-transcribed strand (NTS) of genes can lead to asymmetry in mutation density (*34*). As both SBS19 and SBS5, which are similar to MS_Sig3 and MS_Sig4 respectively, are reported to have transcription strand asymmetry in the COSMIC database (*30*), we investigated this in our data.

Transcription strand asymmetry analysis was performed for all 172 single OL and neuron samples and significant asymmetry was observed for five of six mutation types: C>T (p=1.3×10^−11^), T>C (p<2.0×10^−16^), T>G (p=2.7×10^−13^), C>G (p=8.8×10^−5^), C>A (p=1.9×10^−4^), T>A (p=0.63) (**Fig. 4B**). Stratification of the samples by cell type revealed that the pattern of strand asymmetry is the same for OLs and neurons (**Fig S9**). Log2(TS/NTS) ratios for C>T mutations were generated to allow comparison with those available for similar signatures in COSMIC (*30*) (**Fig. 4C**). Interestingly, three of our signatures showed greater C>T strand asymmetry than for the most similar COSMIC signatures (0.298 for MS_Sig1 vs. 0.070 for SBS44, 0.441 for MS_Sig2 vs. 0.058 for SBS30, 0.324 for MS_Sig4 vs. 0.041 for SBS5), whereas MS_Sig3 (0.445) had lower asymmetry than SBS19 (0.731) (**Fig S8**).

**Fig. 4.**
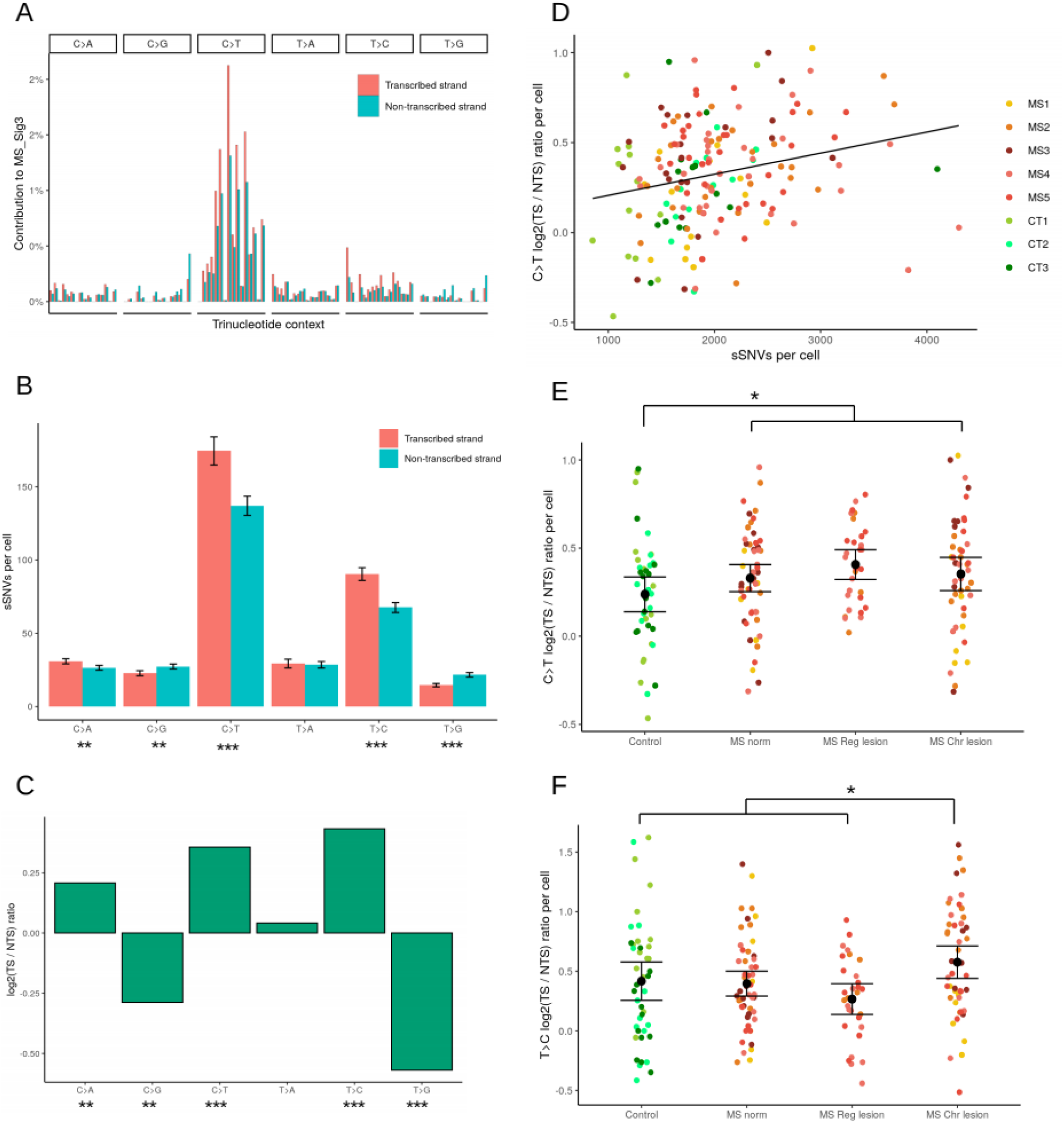
Transcriptional strand asymmetry. **(A)** For MS_Sig3, the mutational signature with strongest transcriptional strand asymmetry, mutation frequencies (x-axis) for six substitution types in trinucleotide context on the template strand (TS) and non-template strand (NTS) of genes. **(B)** Mean sSNVs per cell across all 172 samples for the six mutation types on the TS and NTS with asterisks indicating level of statistical significance for the difference and 95% confidence intervals shown. **(C)** log2(TS/NTS) ratio for each mutation type computed using sSNVs from all 172 samples. **(D)** log2(TS/NTS) ratio for C>T mutations plotted against total SNVs for each cell (coloured circles), indicating a positive correlation. The log2(TS/NTS) ratio from all 172 samples stratified by tissue type for C>T **(E)** and T>C **(F)** mutations, showing associations between TSA and disease status (all MS tissue vs. control) for C>T and MS chronic lesion vs. other tissue types for T>C. Mean values are indicated by a black circle with 95% confidence intervals. Significance levels indicated by asterisks: * p < 0.05, ** p < 10^−3^, *** p < 10^−9^.

We next determined whether different measures: total sSNV count, age, disease status, tissue type and cell type, were associated with transcription strand asymmetry. For C>T mutations, asymmetry was associated with total sSNV count per cell (correlation=0.242, p=0.0019) (**Fig. 4D**), as was disease status (mean log2(TS/NTS) ratio of 0.356 in all MS vs. 0.238 in control tissue, p=0.024) (**Fig. 4E**). Strand asymmetry was less strongly elevated in MS lesion tissue compared to MS normal and control tissue (p=0.083). There were no significant associations for age (p=0.36) or cell type (p=0.29). For T>C, another common mutation exhibiting strong strand asymmetry, significant associations were observed with total sSNVs (p=0.0083) and with chronic MS lesion compared to other tissues (mean log2(TS/NTS) ratio of 0.578 in chronic lesion vs. 0.372 in other tissue, p=0.0061) (**Fig. 4F**), but not for disease status (p=0.90). The only other significant transcription strand asymmetry association was with total sSNVs for C>A substitutions (p=0.027).

## Discussion

For the first time, we report on the mutational landscape of neurons and OLs in MS, the most common acquired chronic neurological disease affecting young adults. This includes the first time, in health or disease, that the mutational landscape of OLs has been reported. A key feature of this work has been the multi-level, controlled study design, permitting direct comparison of single OLs and neurons from the same tissue sample, and differences to be detected between cells from pathologically distinct brain environments within and between individuals. This has enabled us to demonstrate an ∼10 % higher sSNV burden for neurons vs. OLs in pathology-free MS and control brain tissue and, in a subset of MS cases, a similar susceptibility of both cell types to DNA damage when exposed to chronic inflammatory demyelination.

The magnitude of DNA damage observed for lesion-resident cells from MS4, a 73-year old male with primary progressive MS, translated to an increase of 31.6 years in genomic aging. While it is not possible to extrapolate this observation to a direct impact on cellular function and viability, or to know at this stage whether cells from other inflammatory demyelinating lesions in this individual are similarly impacted, it is interesting that this individual had the shortest disease duration of any case we studied and did not received disease modifying immunotherapy (DMI). Lesion-resident cells showing excess sSNVs from another case (MS3) had a shorter, but still considerable, increase in genomic age (15.7 years), and they also did not receive DMI. For the two MS cases who did receive DMI, no sSNV excess was observed in lesion-resident cells, which leads us to speculate on a potential neuroprotective benefit of these treatments.

From a mechanistic perspective, it seems most plausible that increased levels of reactive oxygen species (ROS), a by-product of cellular metabolism, could be responsible for the DNA damage observed to OLs and neurons (*16*). Neurons and OLs are metabolically active cells in a healthy brain, and when injured, metabolic activity increases to repair the damage. It is also plausible that infiltrating peripheral immune cells early in lesion development and pro-inflammatory microglia later on could contribute to a mutagenic environment that damages DNA, thus preventing remyelination and causing neurodegeneration (*3*).

To determine the mutational processes underlying somatic mutagenesis in OLs and neurons, we conducted mutational signature analysis (*27*). A previous study in 161 single neurons identified three signatures (*19*), signature A being associated with aging, which we believe to be equivalent to our MS_Sig4 (SBS5), and two other signatures. Signature B was dominated by C>T transitions, and signature C, which was more balanced with increased C>A transversions. While some of our signatures appeared similar to signature B, none were similar to signature C.

Two of the mutational signatures we discovered, MS_Sig1 and MS_Sig2, had similar signatures in the COSMIC database with proposed aetiologies of defective DNA mismatch repair (SBS44) and defective base excision repair (SBS30), respectively. Thus, while the age-associated signature we identified as MS_Sig4 (SBS5, observed in both health and disease), dominated contributions to sSNV count in all samples, these other signatures, including MS_Sig3 (similar to a signature of unknown aetiology, SBS19, with a high degree of transcriptional strand bias) contributed variably to sSNV counts in samples within and across individuals.

Transcription-associated mutagenesis is an endogenous mutational process that has been shown to occur in single neurons (*18, 35*). Results from our analysis of transcription strand asymmetry, a measure of transcription-associated DNA damage, supports previous findings and suggests it is also a prominent mutagenic mechanism in OLs. We have also demonstrated that aging, disease status and tissue type represent variables that are associated with transcription-associated mutagenesis in our samples. C>T transitions were the most common somatic mutation type we observed, which aligns with findings in previous single neuron studies (*18*). Cytosine deamination is a common biochemical change to DNA that can result in C>T transitions (*36*), however, in the context of gene transcription and our observations of strand asymmetry, namely increased C>T on the transcribed vs. non-transcribed strand, it can be interpreted as increased damage to G on the non-transcribed strand, followed by defective DNA repair. Previous findings of oxidative damage to G in neuronal and OL DNA in MS lesions post-mortem support this observation (*14, 15*).

Here, for the first time, we demonstrate that inflammation in the MS brain is associated with damage to the nuclear genomes of OLs and neurons, albeit in a subset of MS cases. We have demonstrated that the endogenous mutational processes, gene transcription and defective DNA repair are involved. Further work in more cells from more MS brain donors will be required to provide a more comprehensive understanding of the link between neuroinflammation, DNA damage and neurodegeneration. Finally, our observations suggest that therapeutic avenues targeting neuroprotection and DNA repair should be investigated for MS, which could have broader application to other neuroinflammatory conditions and neurodegenerative diseases.

## Acknowledgments

Multiple sclerosis tissue used in this research was provided by the Multiple Sclerosis Research Australia Brain Bank located at the Brain and Mind Centre in Sydney, Australia. The MS Research Australia Brain Bank is supported by Multiple Sclerosis Research Australia, the University of Sydney and the Brain and Mind Centre, the NSW Office for Health and Medical Research, Royal Prince Alfred Hospital Sydney, and has also received support from a number of trusts and foundations including the Trish MS Research Foundation, Levy Foundation, Collier Charitable Fund, Medical Advances Without Animals Trust, and the FIL Foundation. We sincerely thank the MS tissue donors and their families for making this scientific research possible and thank Antony Harding for his assistance. We sincerely thank the control tissue donors and their families, and Fairlie Hinton and Geoff Pavey for assistance with provision of tissue from the Victorian Brain Bank. Our thanks go to Vanta Jameson and Franca Casa Granda from the University of Melbourne (UoM) Flow cytometry core and droplet digital PCR facilities, and also University of Melbourne Research Computing Services. We thank Linda Ngyuen for assistance with figure illustrations. This work received support from China National GeneBank (CNGB).

